# Two cyclic electron flows around photosystem I differentially participate in C_4_ photosynthesis

**DOI:** 10.1101/2022.09.23.509273

**Authors:** Takako Ogawa, Kana Kobayashi, Yukimi Y. Taniguchi, Toshiharu Shikanai, Naoya Nakamura, Akiho Yokota, Yuri N. Munekage

## Abstract

C_4_ plants assimilate CO_2_ more efficiently than C_3_ plants because of their C_4_ cycle that concentrates CO_2_. However, the C_4_ cycle requires additional ATP molecules, which may be supplied by the cyclic electron flow around photosystem I. One cyclic electron flow route, which depends on a chloroplast NADH dehydrogenase-like (NDH) complex, is suggested to be crucial for C_4_ plants despite the low activity in C_3_ plants. The other route depends on proton gradient regulation 5 (PGR5) and PGR5-like photosynthetic phenotype 1 (PGRL1), which is considered a major cyclic electron flow route to generate the proton gradient across the thylakoid membrane in C_3_ plants. However, its contribution to C_4_ photosynthesis is still unclear. In this study, we investigated the contribution of the two cyclic electron flow routes to the NADP-malic enzyme subtype of C_4_ photosynthesis in *Flaveria bidentis*. We observed that the suppression of the NDH-dependent route drastically delayed growth and decreased the CO_2_ assimilation rate to approximately 30% of the wild-type rate. On the other hand, the suppression of the PGR5/PGRL1-dependent route did not affect plant growth and resulted in a CO_2_ assimilation rate that was approximately 80% of the wild-type rate. Our data indicate that the NDH-dependent cyclic electron flow substantially contributes to the NADP-malic enzyme subtype of C_4_ photosynthesis and that the PGR5/PGRL1-dependent route cannot complement the NDH-dependent route in *F. bidentis*. These findings support the fact that during the C_4_ evolution, the photosynthetic electron flow may be optimized to provide the energy required for C_4_ photosynthesis.

## Introduction

Photosynthetic organisms assimilate CO_2_ via the Calvin-Benson cycle using ATP and NADPH produced by photosynthetic electron transport. Linear electron flow (LEF) is driven by photosystem II (PSII) and photosystem I (PSI) and produces both ATP and NADPH. On the other hand, cyclic electron flow (CEF) around PSI generates the proton gradient (ΔpH) across the thylakoid membrane by recycling electrons from the acceptor side of PSI to plastoquinone (PQ) and contributes to the production of only ATP. CEF is important for the regulation of ATP/NADPH production ratio in chloroplasts because ATP/NADPH production by LEF is fixed and insufficient for consumption by CO_2_ assimilation via the Calvin-Benson cycle and photorespiration (Allen, 2003).

CEF has been suggested to be important for C_4_ plants because they require additional ATP to drive CO_2_ concentrating mechanism, called the C_4_ cycle (Hatch, 1987; Kanai and Edwards, 1999; Munekage, 2016). C_4_ plants have evolved from C_3_ plants in multiple lineages in the angiosperm over the past 30 million years (Sage et al., 2012). With the exception of single-cell C_4_ photosynthesis, the C_4_ plants concentrate CO_2_ around ribulose-1,5-bisphosphate carboxylase/oxygenase (RuBisCO) by C_4_ cycle that exchanges four carbon inorganic acids and three carbon inorganic acids between two distinct photosynthetic cells, mesophyll cells (MCs) and bundle sheath cells (BSCs). In the C_4_ cycle, CO_2_ is first converted to HCO_3_^-^ by carbonic anhydrase, which is then fixed to phosphoenolpyruvate to form oxaloacetic acid by phosphoenolpyruvate carboxylase (PEPC) in MCs. The generated oxaloacetic acid is converted to malate or aspartate, which diffuse via plasmodesmata into BSCs, where RuBisCO is located, and decarboxylated to release CO_2_ (Hatch, 1987). Three biochemical subtypes of C_4_ species have been classified based on which of the three enzymes, NADP-malic enzyme (NADP-ME), NAD-malic enzyme (NAD-ME) or phosphoenolpyruvate carboxykinase (PEP-CK), are primarily responsible for C_4_ acids decarboxylation in BSCs (Hatch, 1987; Furbank, 2011). In plants using the C_4_ pathway of the NADP-ME or NAD-ME subtypes, the ATP/NADPH demand for CO_2_ fixation rises to 2.5 because two molecules of ATP are required for regeneration of phosphoenolpyruvate in addition to the three molecules of ATP and two molecules of NADPH required for the Calvin-Benson cycle, whereas in plants with C_3_ photosynthesis, the ATP/NADPH demand ranges between 1.55 and 1.67, depending on photorespiration (Kanai and Edwards, 1999; Osmond, 1981). Furthermore, in the C_4_ pathway of the NADP-ME subtype, NADPH is generated at the step of decarboxylation of malate in BSC. As a result, in the NADP-ME subtype C_4_ species, the ATP/NADPH demand rises in BSC, but in the NAD-ME subtype C_4_ species, it rises in MC. Thus, the balance between LEF and CEF activities differs depending on the subtype and the cell type. Indeed, downregulation of LEF associated with the absence or reduction of grana stacks in chloroplasts of BSC was observed in a number of the NADP-ME subtype C_4_ species, including monocot and eudicot (Andersen et al., 1972; Dengler and Nelson, 1999; Höfer et al., 1992; Woo et al., 1970).

Two CEF routes around PSI have been identified in land plants. One route depends on a chloroplast NADH dehydrogenase-like (NDH) complex, which comprises plastid-encoded subunits (NdhA-K) and more than 19 nuclear-encoded subunits, including NdhL-O specific to photosynthetic NDH (Peltier et al., 2016). Recent studies revealed that the NDH-dependent CEF mediates the transfer of electrons from ferredoxin to PQ (Yamamoto and Shikanai, 2013; Schuller et al., 2019). The other route depends on a proton gradient regulation 5 (PGR5)/PGR5-like photosynthetic phenotype 1 (PGRL1) heterodimer, which is also involved in transferring electrons from ferredoxin to PQ (Munekage et al., 2002; DalCorso et al., 2008; Hertle et al., 2013). These CEF routes are important not only for elevating ATP/NADPH production ratio but also for protecting photosystems from photodamage. The PGR5/PGRL1-dependent route is reported to be involved in ΔpH-dependent regulation of light energy absorption at PSII, detected as nonphotochemical quenching of chlorophyll fluorescence (NPQ). It is also reported that PGR5/PGRL1-dependent CEF is important to prevent PSI over-reduction by limiting electron transport at cytochrome *b_6_f* complexes under strong or fluctuating-light conditions in *Arabidopsis thaliana* and *Oryza sativa* (Munekage et al., 2002, 2004; Suorsa et al., 2012; Yamori et al., 2016). The NDH-dependent route is also believed to act as a safety valve to prevent the stromal over-reduction under stress conditions such as strong light and low temperature (Endo et al., 1999; Yamori et al., 2011), but impairment of NPQ induction or growth defect under fluctuating light conditions is not observed in NDH deficient C_3_ plants except for *Oryza sativa* (Munekage et al., 2004; Yamori et al., 2016; Suorsa et al., 2012). Thus, the PGR5/PGRL1-dependent route is considered to substantially contribute to the CEF activity, whereas the NDH-dependent route is a minor route in C_3_ plants.

Although the NDH-dependent route is less important in C_3_ plants, several reports suggest the importance of the NDH-dependent route in C_4_ plants (Takabayashi et al., 2005; Nakamura et al., 2013). Abundance of NDH subunits was cell-selectively increased, corresponding to the ATP/NADPH demand in some NADP-ME or NAD-ME subtype C_4_ species (Takabayashi et al., 2005; Nakamura et al., 2013). Furthermore, Peterson et al. (2016) showed that the net CO_2_ assimilation rate was decreased by one-half in the transposon insertion lines of *NdhN* or *NdhO* in *Zea mays*, which is an NADP-ME subtype C_4_ species of monocots. Ishikawa et al. (2016) also observed that the *NdhN*-knockdown line of *Flaveria bidentis*, an NADP-ME subtype C_4_ eudicot species, exhibited poor growth under low light conditions, despite relatively mild and largely recovered phenotype under medium light conditions. These reports suggest the importance of the NDH-dependent route for supplying additional ATP in NADP-ME subtype C_4_ species. On the other hand, the extent to which PGR5/PGRL1-dependent ΔpH generation contributes to C_4_ photosynthesis remains unclear. Although abundances of PGR5 and PGRL1 were equal between MC and BSC regardless of the ATP/NADPH demand (Takabayashi et al., 2005; Nakamura et al., 2013), not only the abundance of NDH but also that of PGR5 or PGRL1 was higher in C_4_ species than C_3_ species in genus *Flaveria* (Nakamura et al., 2013), suggesting that the activity of the PGR5/PGRL1-dependent CEF is also enhanced in C_4_ species. Thus, the PGR5/PGRL1-dependent generation of ΔpH is expected to have an equal or greater contribution to supplying additional ATP required for C_4_ photosynthesis than the NDH-dependent CEF.

In this study, we generated *PGR5-, PGRL1*-, and *NdhO*-knockdown *F. bidentis* lines to clarify the contribution of the two CEF routes to C_4_ photosynthesis. In the *NdhO*-knockdown lines, growth was severely delayed at medium to high light intensity, and the net CO_2_ assimilation rate was reduced to 30% compared to the WT plants. On the other hand, in the *PGR5*- or *PGRL1*-knockdown lines, growth was normal, but the net CO_2_ assimilation rate was reduced to 80% of the WT plants at high light intensity. From the comparative analysis of these lines, we concluded that the NDH-dependent CEF contributes as the major route for C_4_ photosynthesis, and the PGR5/PGRL1-dependent CEF also partly contributes to C_4_ photosynthesis under high light, but the physiological importance of this route is more inclined to the NPQ induction in *F. bidentis.*

## Results

We knocked down *PGR5, PGRL1*, and *NdhO* in *F. bidentis* via RNA interference (RNAi). The target transcript levels in the transgenic plants, in which *PGR5, PGRL1*, or *NdhO* were knocked down (*PGR5*-RNAi, *PGRL1*-RNAi, and *NdhO*-RNAi, respectively), were approximately 10% of the WT levels (Fig. 1A). The expression of all three *PGR5* genes identified in the *Flaveria* genome (*PGR5A, PGR5B*, and *PGR5C*; Taniguchi et al., 2021) was suppressed in the *PGR5*-RNAi lines. The protein levels of PGR5, PGRL1, and NdhH were undetectable or less than 6% of the WT levels in the *PGR5*-RNAi, *PGRL1*-RNAi, and *NdhO*-RNAi lines, respectively (Fig. 1B). Moreover, PGR5 was undetectable in the *PGRL1*-RNAi lines, likely because PGR5 is anchored to the thylakoid membrane by PGRL1 (Hertle et al., 2013). The PGRL1 content in the *PGR5*-RNAi plants was less than 30% of the WT level (Fig. 1, B and C; Supplemental Fig. S1). This may be due to reduced transcription of *PGRL1* (Fig. 1A) and the impaired stability of the PGR5/PGRL1 heterodimer, which has not been observed in the *A. thaliana pgr5* mutant (Hertle et al., 2013). In *PGR5-* and *PGRL1*-RNAi plants, the amounts of NdhH and Rieske, a subunit of cytochrome *b_6_f* complex, were similar to the corresponding amounts in WT plants. In *NdhO*-RNAi plants, the amounts of PGR5 and PsaD, a subunit of PS I, were similar to those in WT plants, while amounts of PGRL1, PsbO, a subunit of PS II, and Reiske tended to decrease, although the decrease was not statistically significant, compared to vector control when the amounts of WT plants were used as the standard (Fig. 1, B and C; Supplemental Fig. S1).

**Figure 1.**
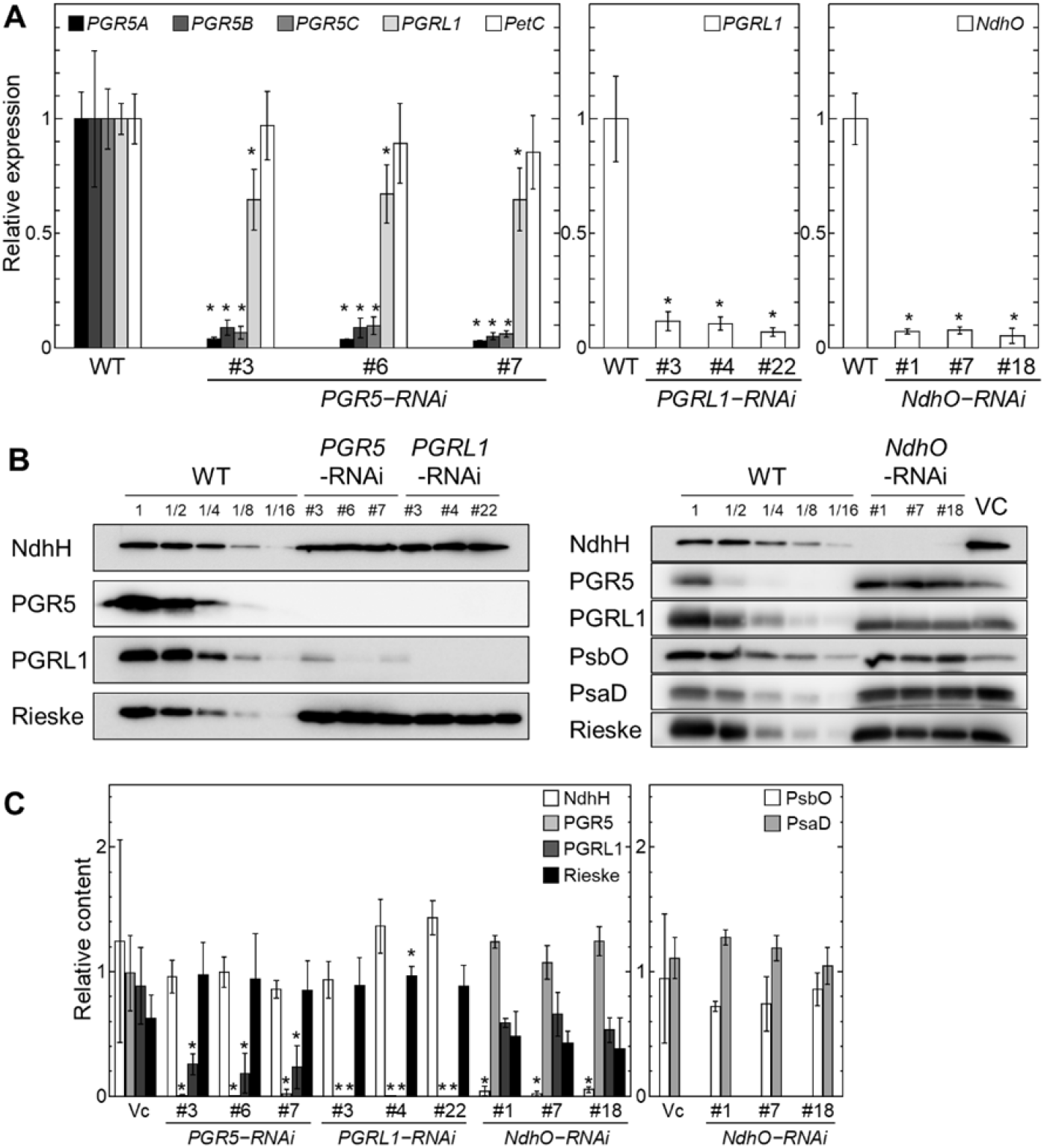
Knockdown of *PGR5, PGRL1,* and *NdhO* in *F. bidentis.* (A) Expression of *PGR5A, PGR5B, PGR5C,* and *PGRL1* in *PGR5*-RNAi lines, *PGRL1* in *PGRL1*-RNAi lines, and *NdhO* in *NdhO*-RNAi lines relative to the corresponding expression in the WT controls. Vertical bars indicate the SD (n = 4–5). Asterisks indicate significant differences (Student *t*-test, *P* < 0.05) between WT and *PGR5*-RNAi, *PGRL1*-RNAi, or *NdhO*-RNAi lines. (B) Immunoblot analysis of the membrane proteins extracted from the leaves of the WT, vector control (VC), *PGR5*-RNAi, *PGRL1*-RNAi, and *NdhO*-RNAi lines. Lanes were loaded with 20 μg protein to detect PGR5 and 10 μg to detect PGRL1, NdhH, Rieske, PsaD and PsbO. The dilution series for the WT plants are indicated. (C) Relative content of membrane proteins involved in cyclic or linear electron flow. The amount of proteins was quantified by chemiluminescence signal intensities of immunoblot analysis, and the signal intensity of the WT plants was set to 1. Vertical bars indicate the SD (n=3). Asterisks indicate significant differences (Student *t*-test, *P* < 0.05) between VC and *PGR5*-RNAi, *PGRL1*-RNAi, or *NdhO*-RNAi lines.

To assess CEF activities in the RNAi lines, we monitored the ferredoxin-dependent transfer of electrons to PQ, which was reflected by an increase in chlorophyll fluorescence following the addition of NADPH and ferredoxin to ruptured chloroplasts (Fig. 2). Ruptured chloroplasts from homogenized leaves contained both BSC and MC chloroplasts, as judged by the presence of PEPCs localized in MCs and RbcL, a large subunit of RuBisCO, localized in BSCs in the homogenized leaf supernatant (Supplemental Fig. S2). PQ reduction level was estimated with time-dependent chlorophyll fluorescence level (Ft), normalized as (Ft-Fo)/(Fm-Fo), in which Fo was chlorophyll fluorescence level before the addition of NADPH and ferredoxin, where Q_A_ was oxidized, and Fm was that gained by illumination with saturating pulse, where Q_A_ was fully reduced. The increase in chlorophyll fluorescence level after the addition of ferredoxin was delayed in the RNAi lines compared with the WT plants (Fig. 2), suggesting the suppression of ferredoxin-dependent electron transfer to PQ via CEF in ruptured chloroplasts of the RNAi lines. The application of antimycin A, which inhibits the PGR5/PGRL1-dependent route (Munekage et al., 2002), delayed the increase in chlorophyll fluorescence in WT plants. Ferredoxin-dependent PQ reduction was not delayed by antimycin A in *PGR5*-RNAi and *PGRL1*-RNAi plants, whereas it was considerably delayed in *NdhO*-RNAi plants. This indicates that there are two independent cyclic routes in *F. bidentis*; antimycin A sensitive route was suppressed in *PGR5*- and *PGRL1*-RNAi plants, and antimycin A insensitive route was suppressed in *NdhO*-RNAi plants. The final level of chlorophyll fluorescence was elevated in the presence of antimycin A in WT, *PGR5*-RNAi, and *PGRL1*-RNAi plants. This could be a secondary effect of antimycin A, which may inhibit electron transport downstream of PQ.

**Figure 2.**
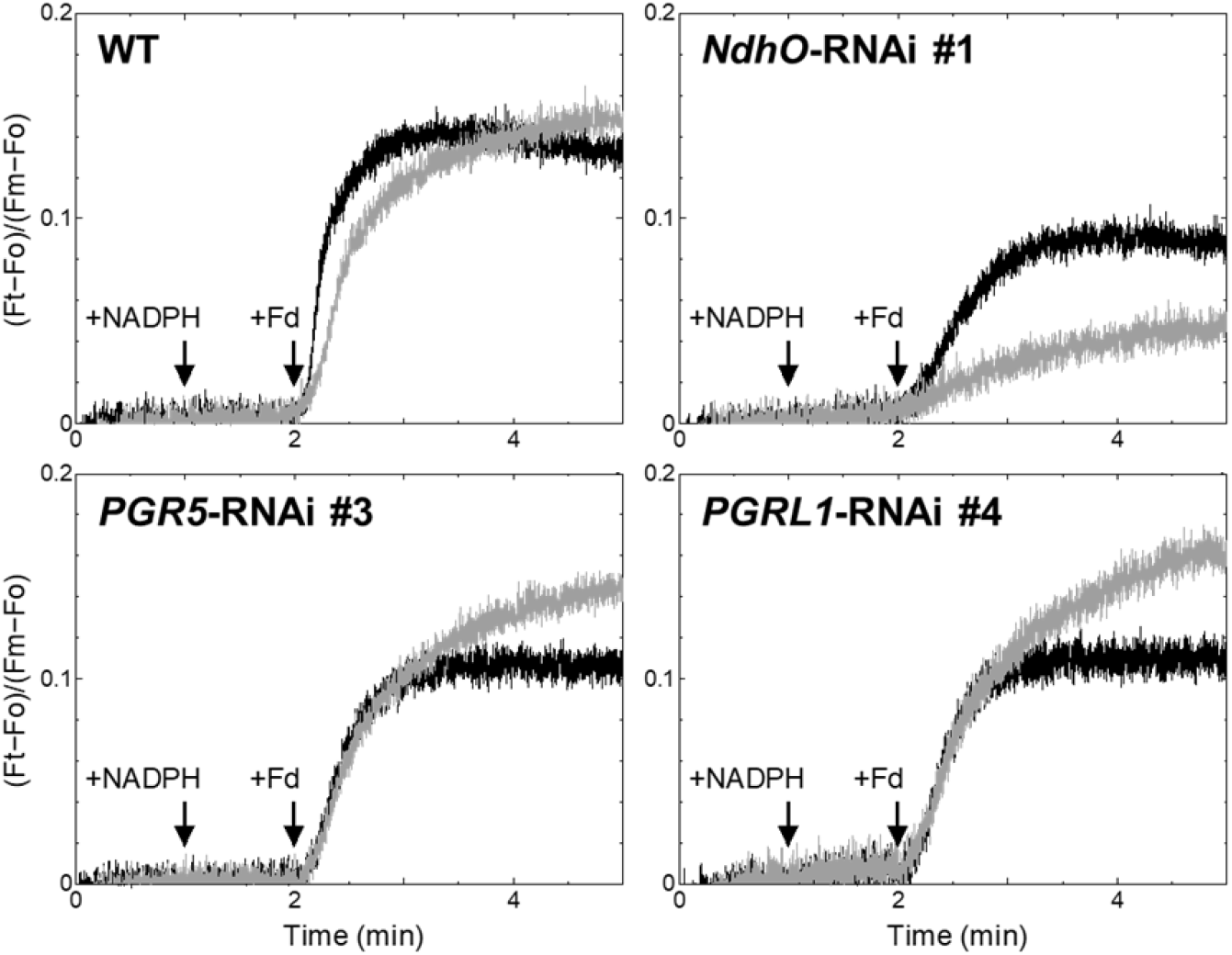
Electron transfer to PQ in ruptured chloroplasts (20 μg chlorophyll ml^-1^) following the addition of 250 μM NADPH and 5 μM ferredoxin (Fd). The electron transfer was based on the chlorophyll fluorescence under weak light (0.25 μmol photons m^-2^ s^-1^). Data are presented as the average of three measurements. Black line, no antimycin A; gray line, in the presence of 5 μM antimycin A.

The ferredoxin-dependent PQ reduction level in ruptured chloroplasts was lower in CEF-suppressed plants than in WT plants, whereas the maximum quantum yield of PSII (Fv/Fm) and the effective quantum yield of PSII (Φ_PSII_) in the ruptured chloroplasts treated with methyl viologen, an electron acceptor of PSI, of the *PGR5*-RNAi and *PGRL1*-RNAi plants were similar to the corresponding values in WT plants (Table 1). These findings suggest that LEF activities were not impaired in the ruptured chloroplasts of *PGR5*-RNAi and *PGRL1*-RNAi plants. However, Fv/Fm and Φ_PSII_ in the ruptured chloroplasts were lower in *NdhO*-RNAi plants than in WT plants, suggesting that the LEF activity was impaired in *NdhO*-RNAi plants.

**Table 1.** Quantum yield of ruptured chloroplasts in the presence of methyl viologen.

| | Fv/Fm | $\Phi_{PSII}$ (260 $\mu\text{mol photons m}^{-2} \text{s}^{-1}$) |
| --- | --- | --- |
| Wild-type | 0.765 ± 0.012 | 0.302 ± 0.031 |
| *PGR5*-RNAi #3 | 0.765 ± 0.004 | 0.244 ± 0.034 |
| *PGRL1*-RNAi #4 | 0.777 ± 0.003 | 0.322 ± 0.031 |
| *NdhO*-RNAi #1 | 0.706 ± 0.008* | 0.178 ± 0.016* |

Data are presented as the average ± SD (n = 3–4). The asterisk indicates a significant difference (Student *t*-test, *P* < 0.05) between the wild-type plants and the *PGR5*-RNAi, *PGRL1*-RNAi, or *NdhO*-RNAi plants.

The *NdhO*-RNAi plants were smaller than the WT plants grown under 250 μmol photons m^-2^ s^-1^ illumination for 45 days (Fig. 3A). The leaf area of *NdhO*-RNAi lines decreased to 3–7% of that of WT plants (Fig. 3B). Furthermore, *NdhO*-RNAi plants took twice as long as WT plants to flower (Fig. 3C), although the *NdhO*-RNAi and WT plants both set the first flower bud at the 12^th^ node. Even under 1,000 μmol photons m^-2^ s^-1^, the *NdhO*-RNAi plants were smaller than the WT plants and the leaf area was decreased to 10–14% compared to that of the WT plants (Supplemental Fig. S3). These observations indicate drastically slow growth of the *NdhO*-RNAi plants in comparison with the WT plants. In contrast, the growth of *PGR5*-RNAi and *PGRL1*-RNAi plants was similar to that of WT plants under 250 μmol photons m^-2^ s^-1^ and 1,000 μmol photons m^-2^ s^-1^ (Fig. 3 and Supplemental Fig. S3). Thus, the NDH-dependent CEF route, but not the PGR5/PGRL1-dependent route, is crucial for normal plant growth under both medium and high light conditions.

**Figure 3.**
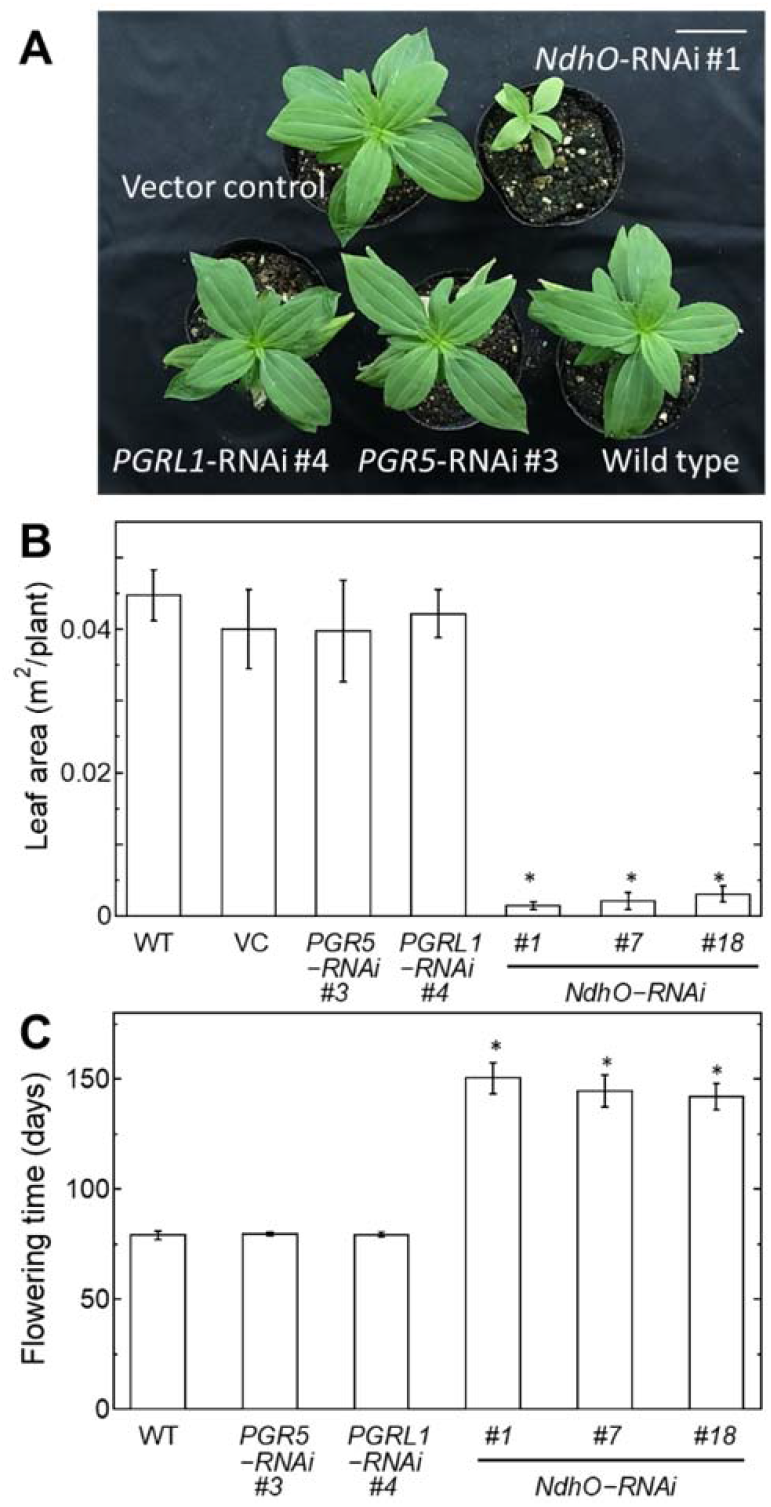
Photosynthetic activity assessed based on growth under medium light condition (250 μmol photons m^-2^ s^-1^). (A) Observable phenotypes of 45-day-old plants. The bar indicates 5 cm. (B) Leaf area per plant of 45-day-old plants. (C) Days to flowering. Vertical bars indicate the SD (n = 5). Asterisks indicate significant differences (Student *t*-test, *P* < 0.05) between WT and *PGR5*-RNAi, *PGRL1-RNAi,* or *NdhO*-RNAi lines.

The net CO_2_ assimilation rate of *NdhO*-RNAi #1 and #18 decreased to 28% and 46% of that of WT plants, respectively, at 2,000 μmol photons m^-2^ s^-1^ (Fig. 4A). Considering the chlorophyll content per leaf area in *NdhO*-RNAi #1 and #18 was 55% and 66% of that in WT plants, respectively (Table 2), the net CO_2_ assimilation rate per chlorophyll content in *NdhO*-RNAi #1 and #18 decreased to approximately 50% and 70% of that in WT plants, respectively. Compared with the effects of the knockdown of *NdhO*, the knockdown of *PGR5* and *PGRL1* had less of an impact on the net CO_2_ assimilation rate. However, the net CO_2_ assimilation rate per leaf area in *PGR5*-RNAi and *PGRL1*-RNAi plants decreased to approximately 80% of that in WT plants at 2,000 μmol photons m^-2^ s^-1^ (Student *t*-test, *P* < 0.05, Fig. 4A). The chlorophyll content per leaf area in *PGR5*-RNAi and *PGRL1*-RNAi plants did not decrease compared with the WT plants (Table 2). Therefore, we conclude that the NDH-dependent CEF route substantially contributes to the CO_2_ assimilation in *F. bidentis*, whereas the PGR5/PGRL1-dependent route makes a small contribution to the CO_2_ assimilation.

**Figure 4.**
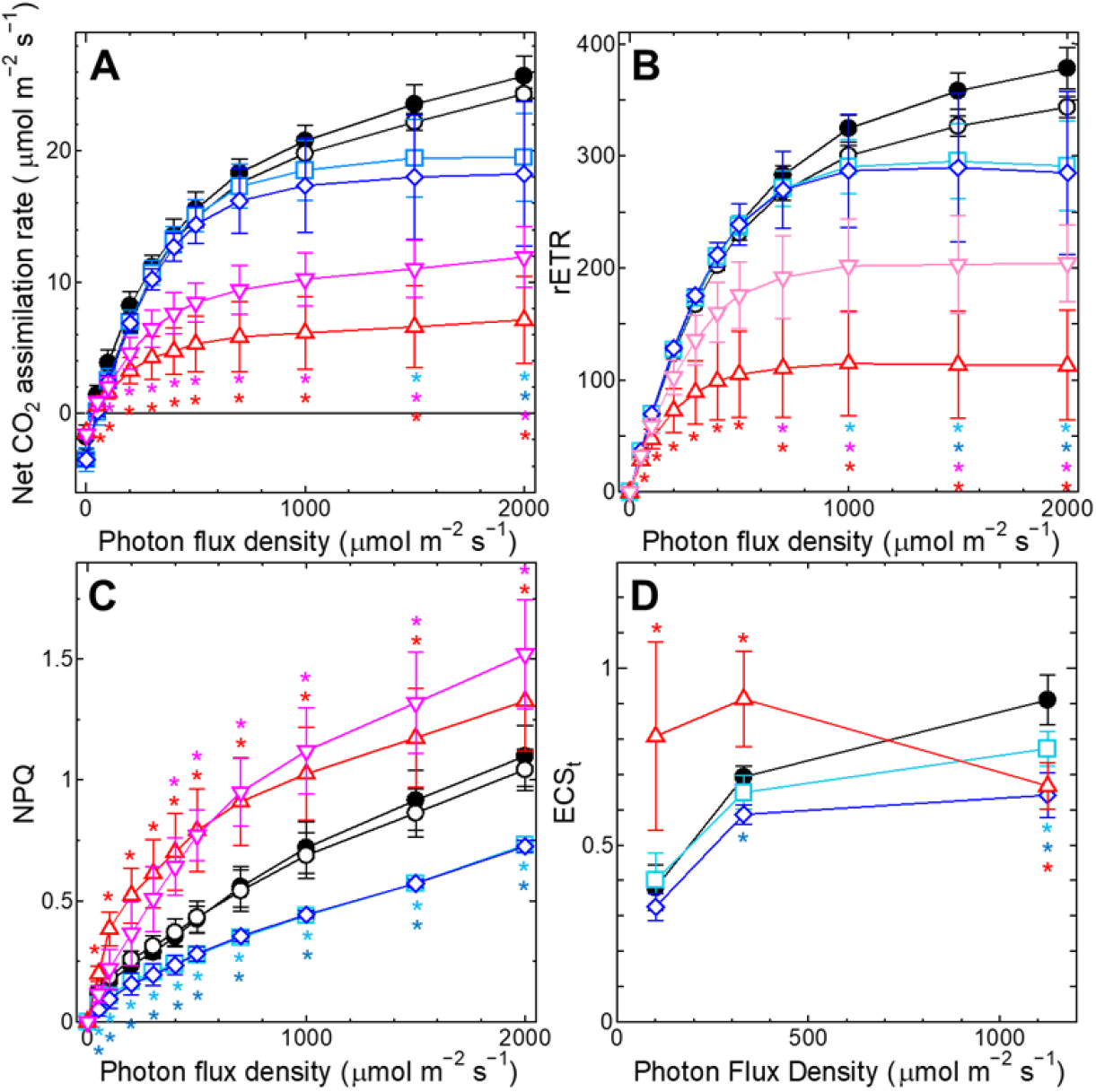
Response curve of the net CO_2_ assimilation rate per leaf area (A), rETR (B), NPQ (C) and the ECSt parameter (D) to light intensity. (A-C) Chlorophyll fluorescence parameters were measured along with the CO_2_ assimilation rate under 400 ppm ambient CO_2_. (D) The ECSt parameter was estimated by measuring the rapid decline of the electrochromic pigment absorbance shift after cessation of actinic light. Black closed circles, WT; black open circles, vector control; light blue squares, *PGR5*-RNAi #3; blue diamonds, *PGRL1*-RNAi #4; red triangles, *NdhO*-RNAi #1; pink inverted triangles, *NdhO*-RNAi #18. Vertical bars indicate the SD (A-C, n = –6; D, n = 4–6). Light blue, blue, red or pink asterisks indicate significant differences (Student *t*-test, *P* < 0.05) between WT and *PGR5*-RNAi #3, *PGRL1-RNAi* #4, *NdhO*-RNAi #1 or *NdhO*-RNAi #18, respectively.

**Table 2.** Chlorophyll content per leaf area.

| | Chlorophyll *a* + *b* ($\mu\text{g mm}^{-2}$) |
| --- | --- |
| Wild-type | 0.47 ± 0.03 |
| *PGR5*-RNAi #3 | 0.52 ± 0.04 |
| *PGRL1*-RNAi #4 | 0.51 ± 0.02* |
| *NdhO*-RNAi #1 | 0.26 ± 0.05* |
| *NdhO*-RNAi #18 | 0.31 ± 0.02* |

Data are presented as the average ± SD (n = 5). The asterisk indicates a significant difference (Student *t*-test, *P* < 0.05) between the wild-type plants and the *PGR5*-RNAi, *PGRL1*-RNAi, or *NdhO*-RNAi plants.

Decreased net CO_2_ assimilation rate in the transgenic lines (Fig. 4A) may be caused by decreased ΔpH due to the suppressed CEF activity. To confirm the electron transport activity and ΔpH, we measured chlorophyll fluorescence parameters (Fig. 4, B and C) along with the CO_2_ assimilation rate (Fig. 4A). Notably, chlorophyll fluorescence detected here was mainly derived from chloroplasts in the first MC layer. In C_4_ plants, the PSII content is low in the BSC (2–23%) compared with that in the MC (Höfer et al., 1992), so the chlorophyll fluorescence parameters mainly reflect the electron transport in MCs. Thus, the net CO_2_ assimilation rate (Fig. 4A) reflects the results of coordinated photosynthetic activities of MC and BSC, while chlorophyll fluorescence parameters (Fig. 4, B and C) reflect the photosynthetic activity of only MC. The rETR, which is the chlorophyll fluorescence parameter representing the relative electron transfer rate of LEF, was calculated as the product of Φ_PSII_ and photosynthetically active radiation. The rETR decreased in parallel with the net CO_2_ assimilation rate in the CEF-suppressed plants (Fig. 4, A and B; Supplemental Fig. S4). Since Φ_PSII_ in ruptured chloroplasts of *PGR5*-RNAi and *PGRL1*-RNAi plants in the presence of methyl viologen was similar to that in WT plants (Table 1), indicating that linear electron transport itself is not affected, the restricted rETR in MC of the leaves of *PGR5*-RNAi and *PGRL1*-RNAi plants (Fig. 4B) is likely due to the decrease in CO_2_ assimilation (Fig. 4A). We estimated ΔpH by the magnitude of NPQ because NPQ mainly reflects the thermal dissipation triggered by lumen acidification in land plants (Ruban, 2016). While the NPQ of *PGR5*-RNAi and *PGRL1*-RNAi plants was lower than that of WT plants, it increased in *NdhO*-RNAi plants (Fig. 4C). We also analyzed the size of proton motive force (*pmf*) in the CEF-suppressed plants by measuring the rapid decline of the electrochromic pigment absorbance shift (ECS) when the low, medium, or high light (100, 330, or 1130 μmol m^-2^ s^-1^, respectively) is switched off (Fig. 4D; Supplementary Fig. S7). The ECSt parameter reflects the light–dark difference in the membrane potential formed across the thylakoid membrane (Cruz et al., 2004). ECS signal is primarily based on carotenoids and chlorophyll *b*, which are mainly associated with PSII. Since this system detects the change in absorbance of light transmitted through the leaf, the ECS signal is derived from chloroplasts in both MC and BSC. In *F. bidentis*, however, since chloroplasts in BSC contain less PSII, the ECS signal is expected to be highly dependent on chloroplasts in MC. In the WT plants, the ECSt rose with the increase in light intensity (Fig. 4D). The ECSt in *PGRL1*-RNAi plants was similar to that in WT plants under low light but was lower under medium and high light. In contrast, the ECSt in *NdhO*-RNAi plants was higher than that of WT plants under low and medium light. *NdhO*-RNAi plants exhibited both mild and severe phenotypes in the ECS measurements (Fig. 4D, n=6 for *NdhO*-RNAi #1), leading to a larger standard deviation, possibly because the rate of carbon dioxide fixation was severely reduced in *NdhO*-RNAi plants (Fig 4A), and this metabolic rate differed among individuals. Results of ECSt were consistent with those of light intensity-dependent NPQ induction except for the values for *NdhO*-RNAi plants at high light (Fig. 4C). The decrease in ECSt of the *NdhO*-RNAi plants under high light could be attributed to photodamage caused by prolonged high light exposure during the measurement. The *pmf* is more dependent on the ΔpH than the membrane potential at high light intensity (Cruz et al., 2004). Thus, these results showed that in *F. bidentis*, the PGR5/PGRL1-dependent CEF route contributes to the generation of ΔpH, which is required for NPQ induction in MCs, especially under high light condition, while the NDH-dependent route may be important for the generation of ΔpH in BSC but does not contribute to NPQ induction in MC. *NdhO*-RNAi plants showed higher NPQ and *pmf* than WT plants at a low and medium light intensity, possibly due to slower ATP consumption in MC chloroplasts caused by reduced photosynthetic activity. This could be the result of ATP depletion in BSCs due to the loss of NDH-dependent CEF activity.

To investigate whether the CO_2_ concentration mechanism is affected in the CEF-suppressed plants, we measured the intercellular CO_2_ concentration-dependency of the net CO_2_ assimilation rate (Fig. 5A). The initial slope of the net CO_2_ assimilation rate response curve to intercellular CO_2_ concentration (μmol m^-2^ s^-1^ ppm^-1^) was significantly lower in *PGR5-* or *PGRL1*-RNAi plants (0.11 ± 0.04 or 0.16 ± 0.07, respectively; Student *t*-test, *P* < 0.05) compared to those in wild-type plants (0.27 ± 0.06), and much lower in *NdhO*-RNAi #1 or #18 (0.02 ± 0.02 or 0.07 ± 0.03, respectively; Student *t*-test,*P* < 0.05). These results suggest that the CO_2_ concentration mechanism might be impaired in the CEF-suppressed plants. The maximum net CO_2_ assimilation rate under high CO_2_ concentration was decreased to 85% in *PGR5*-RNAi or *PGRL1*-RNAi plants and 30–50% in *NdhO*-RNAi plants compared to that of WT plants (Fig. 5A). Taking into account the chlorophyll content per leaf area (Table 2), the initial slope of the CO_2_ response curve (μmol gChl^-1^ s^-1^ ppm^-1^) was significantly lower in the CEF-suppressed plants (0.21 ± 0.08, 0.32 ± 0.14, 0.09 ± 0.09, and 0.22 ± 0.11 in *PGR5*-RNAi, *PGRL1*-RNAi, *NdhO*-RNAi #1, and *NdhO*-RNAi #18, respectively; Student *t*-test, *P* <0.05) than in the WT plants (0.58 ± 0.12), whereas the maximum CO_2_ assimilation rate under high CO_2_ concentration was decreased to 80% in *PGR5*-RNAi or *PGRL1* -RNAi plants or 52–75% in *NdhO*-RNAi plants compared to that in WT plants. Since activities of PEPC and RuBisCO could affect the initial slope and saturation rate of the CO_2_ response curve, respectively (von Caemmerer and Furbank, 1999), we examined the amount of PEPC and RbcL in the leaves per chlorophyll basis and found that they neither reduced in *NdhO*-RNAi plants nor in *PGR5*-RNAi or *PGRL1*-RNAi plants compared with that in WT plants (Supplemental Fig. S5). Thus, the lowered maximum CO_2_ assimilation rate and the initial slope of the curve (Fig. 5A) cannot be explained by PEPC or Rubisco activity, suggesting that CO_2_ assimilation rate in these CEF-suppressed lines was limited by reduced activity of phosphopyruvate (PEP) regeneration or Calvin-Benson cycle. Further, we found that chloroplasts, gathered centripetally toward the vein in BSCs of WT, *PGR5*-RNAi, and *PGRL1*-RNAi plants, were scattered in the BSCs of the *NdhO*-RNAi plants (Fig. 5B). Since the centripetal arrangement of chloroplasts in the BSCs is thought to minimize CO_2_ leakage to the MCs, the abnormal position of chloroplasts in the BSCs in *NdhO*-RNAi plants may cause a defect in the CO_2_-concentration mechanism (von Caemmerer and Furbank, 2003).

**Figure 5.**
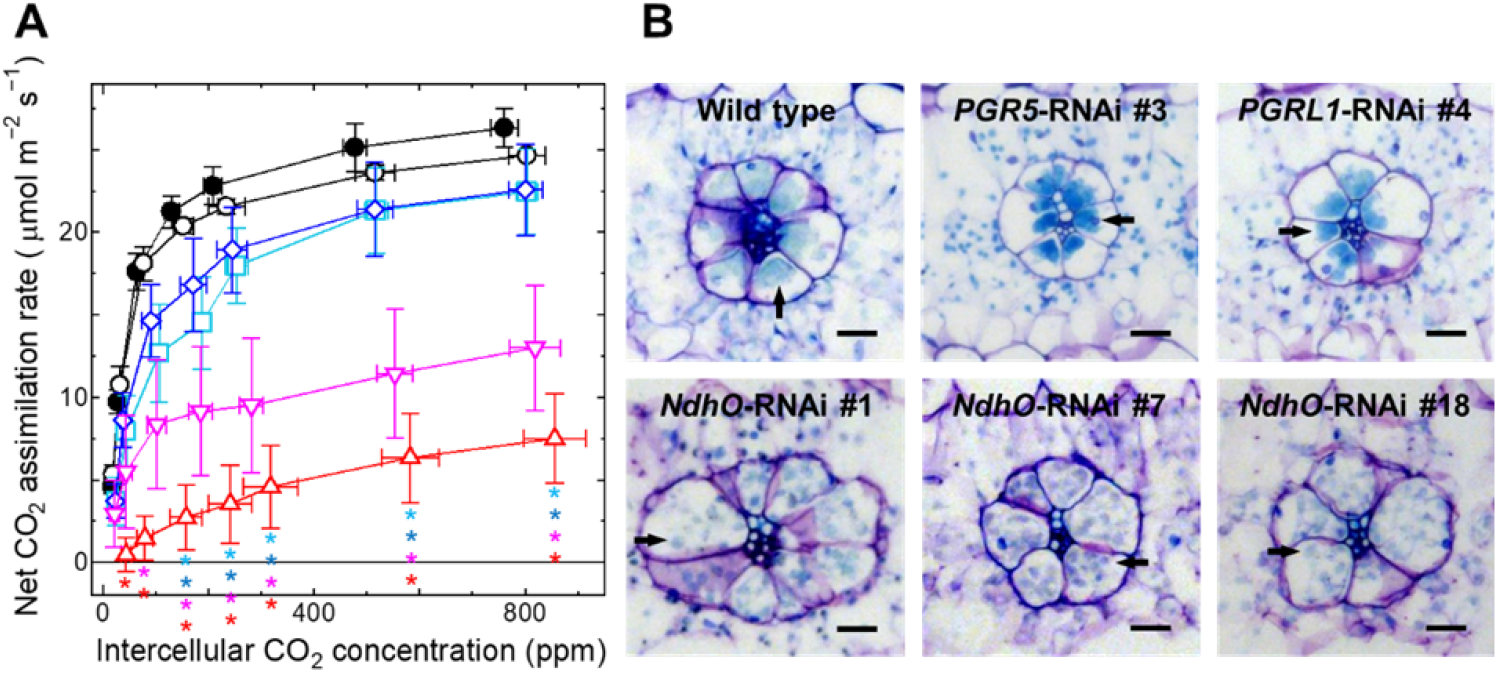
Response curve of the net CO_2_ assimilation rate per leaf area to intercellular CO_2_ concentration under 1,500 μmol m^-2^ s^-1^ illumination (A) and location of chloroplasts in the cell (B). (A) Black closed circles, WT; black open circles, vector control; light blue squares, *PGR5*-RNAi #3; blue diamonds, *PGRL1-RNAi* #4; red triangles, *NdhO*-RNAi #1; pink inverted triangles, *NdhO*-RNAi #18. Vertical bars indicate the SD (n = 3–7). Light blue, blue, red or pink asterisks indicate significant differences (Student *í*-test, *P* < 0.05) between WT and *PGR5*-RNAi #3, *PGRL1-RNAi* #4, *NdhO*-RNAi #1 or *NdhO*-RNAi #18, respectively. (B) Cross section of WT and RNAi plant leaves stained with toluidine blue. Arrows indicate chloroplasts in BSC. Bars indicate 20 μm.

We concluded that impaired CEF results in insufficient ATP production, which limits metabolism, such as PEP regeneration and the Calvin-Benson cycle.

## Discussion

Based on a comparison of *PGR5*-RNAi, *PGRL1*-RNAi, and *NdhO*-RNAi plants, we proved that the NDH-dependent CEF contributes majorly and PGR5/PGRL1-dependent CEF contributes partly to the NADP-ME subtype C_4_ photosynthesis in *F. bidentis*. Although the PGR5/PGRL1-dependent route plays a major role in ΔpH formation in C_3_ plants (Yamori and Shikanai, 2016), the present study showed that the NDH-dependent route plays a central role in the NADP-ME subtype C_4_ photosynthesis in *F. bidentis*.

In C_3_ plants, disruption of the NDH-dependent route had little effect on photosynthesis and plant phenotype, except under stress conditions or in the case of the absence of the PGR5/PGRL1-dependent route, as observed in the double knockout plants of PGR5/PGRL1-dependent and NDH-dependent routes (Endo et al., 1999; Suorsa et al., 2012; Munekage et al., 2004). The NDH complex forms a supercomplex with PSI, but it is only 1–4% of the total PSI in *A. thaliana* (Yamori and Shikanai, 2016; Pribil et al., 2014). Few species have lost the NDH complex, such as *Pinus thunbergii* and *Phalaenopsis aphrodite*, indicating that ATP/NADPH can be optimized without NDH activity in these species (Yamori and Shikanai, 2016). On the other hand, NDH complexes are abundant in C_4_ species and accumulate in a cell-selective manner consistent with the demand for ATP/NADPH, while PGR5 and PGRL1 are equally distributed in MC and BSC (Takabayashi et al., 2005; Nakamura et al., 2013). In the genus *Flaveria*, a subunit of NDH was ten times more abundant in C_4_ species than in C_3_ species (Nakamura et al., 2013). Furthermore, the NdhH amount and NDH activity were increased in C_3_–C_4_ intermediate *Flaveria ramosissima* and C_4_-like *Flaveria brownii*, but the amounts of PGR5 and PGRL1 were not increased (Nakamura et al., 2013; Munekage and Taniguchi, 2016). These evidences suggested that the NDH-dependent route was primarily used to satisfy the ATP/NADPH demand, which increased along with the development of the C_4_ cycle-dependent CO_2_ concentration system during C_4_ evolution. We showed here that impairment of the NDH-dependent route alone resulted in significant growth reduction and decreased photosynthesis in *F. bidentis* (Fig. 3, 4A). Furthermore, we found abnormal chloroplast position in the BSC of the *NdhO*-RNAi plants (Fig. 5B). In C_4_ species that do not have a thick cell wall or a suberin layer between the BSC and MC, chloroplasts are often located at the centripetal position of the BSC (Edwards, 2011). These chloroplasts are distanced from the cell surface on the MC side by vacuoles, suggested to minimize CO_2_ leakage from the BSC to the MC (von Caemmerer and Furbank, 2003). Thus, in *NdhO*-RNAi plants, the abnormal chloroplast position may impair the CO_2_ concentrating mechanism, as reflected in the low initial slope of the net CO_2_ assimilation rate response curve to intercellular CO2 concentration (Fig. 5A).

Previously, Ishikawa et al. (2016) reported that in *NdhN*-knocked-down *F. bidentis* grown at 400 μmol photons m^-2^ s^-1^, net CO_2_ assimilation rates were slightly lower than wild-type plants at light intensities below 1000 μmol photons m^-2^ s^-1^, but were not affected at higher light intensities. However, we observed greater phenotypic differences between *NdhO*-RNAi and WT plants grown both at 250 and 1,000 μmol photons m^-2^ s^-1^ and a drastic decrease of the net CO_2_ assimilation rate in *NdhO*-RNAi plants under high light conditions as well as under low light condition (Figs 3 and 4A; Supplemental Fig. S3), suggesting a larger role of the NDH-dependent CEF in the C_4_ photosynthesis of *F. bidentis.* This discrepancy between the report by Ishikawa et al. (2016) and our results is likely due to the remaining expression and activity of the NDH complex in the *NdhN*-knocked-down lines. A large contribution of the NDH-dependent CEF to C_4_ photosynthesis was also observed in *Z. mays*; knocking out *NdhN* or *NdhO* decreases the CO_2_ assimilation rate by 50% (Peterson et al., 2016). Although monocot *Z. mays* and eudicot *F. bidentis* are phylogenetically distant, interestingly, CEF activity via the NDH complex plays an important role in both these species that have acquired NADP-ME subtype C_4_ metabolic pathways.

It is unclear why the NDH-dependent route, rather than the PGR5/PGRL1-dependent route, became the main route used in the NADP-ME subtype of C_4_ photosynthesis. One possible explanation is that the NDH complex itself functions as a proton pump and thus the NDH-dependent route can generate ΔpH more efficiently than the PGR5/PGRL1-dependent route (Shikanai, 2007). Structural studies have revealed that cyanobacterial NDH-1MS (NDH-I_3_) has proton channels across the thylakoid membrane and a ferredoxin binding site (Schuller et al., 2020; Pan et al., 2020). Since CEF through the NDH complex allows proton translocation across the thylakoid membrane during electron transfer from ferredoxin to PQ in addition to proton translocation by returning electrons at the cytochrome *b_6_f* complex, it may be suitable for the formation of ΔpH to drive ATP synthesis under conditions where electron input from PSII is limited, such as in chloroplasts of BSC.

We showed here that the PGR5/PGRL1-dependent CEF partly helps to supply extra ATP for C_4_ photosynthesis of *F. bidentis* at a high light intensity, based on the result of reduced net CO_2_ assimilation rate in the *PGR5*-RNAi and *PGRL1*-RNAi plants above a light intensity of 1000 μmol photons m^-2^ s^-1^ (Fig 4A). PGR5 and PGRL1 function in the same CEF route in *F. bidentis* as previously shown in *A. thaliana* (Hertle et al., 2013) because we did not see any phenotypic difference between *PGR5*-RNAi and *PGRL1*-RNAi lines in growth, photosynthetic activity, and NPQ inductions (Figs 2,3 and 4), and the absence of PGRL1 led to destabilization of PGR5 and vice versa in C_4_ *F. bidentis* (Fig. 1B). The *PGR5*-RNAi and *PGRL1*-RNAi plants exhibited lower NPQ induction and ECSt than WT plants under high light condition (Fig. 4, C and D), suggesting that the PGR5/PGRL1-dependent route helps generate the ΔpH in *F. bidentis*. However, the PGR5/PGRL1-dependent route cannot complement the NDH-dependent route concerning the production of ATP for C_4_ photosynthesis. This may be because its physiological role is different from that of the NDH-dependent route; the PGR5/PGRL1-dependent CEF may be active and generate ΔpH for NPQ induction at medium to high light intensity, i.e., under the condition of sufficient or excessive light energy, but are not active at a low light intensity with deficient light energy. In contrast, the NDH-dependent CEF may function in ΔpH generation to produce ATP in a broad range of light intensity. This idea was supported by the evidence that lack of PGR5-dependent CEF did not influence *pmf* generation at low light; however, lack of NDH-dependent CEF slightly but significantly affected *pmf* generation at all light intensities in *A. thaliana* (Wang et al., 2015). Moreover, it has been reported that overexpression of PGR5 enhanced an electron sink downstream of PSI but did not increase *pmf* or CO_2_ assimilation rate in *F. bidentis* (Tazoe et al., 2020), suggesting that PGR5/PGRL1-dependent CEF plays a role in avoiding over-reduction of PSI acceptor side and in photoprotection of PSI. Therefore, the PGR5/PGRL1-dependent CEF may be important for photosynthetic regulation rather than ATP synthesis.

C_4_ photosynthesis operates different parts of the metabolic pathway in MCs and BSCs; thus, it is necessary to change the balance between LEF and CEF activities according to ATP/NADPH demand for each cell. In the pure NADP-ME subtype of C_4_ photosynthesis, in which all oxaloacetate produced by PEP carboxylation is converted to malate, it has been assumed that ATP is produced only by CEF activity since NADPH is produced by malate decarboxylation in BSC (Kanai and Edwards, 1999). In *Sorghum bicolor*, presumed to be a pure NADP-ME subtype species, and *Z. mays*, though reported to possess PEP-CK activity (Wingler et al., 1999; Furbank, 2011), LEF activity was almost completely downregulated by repression of PSII subunit expression (Woo et al.,1970; Andersen et al., 1972). In C_4_ *Flaveria* species, because 35–40% of oxaloacetate is converted to aspartate in MC (Moore and Edwards, 1986), which is exported to the BSC where it is converted back to oxaloacetate and reduced to malate by using NADPH, the remaining activity of PSII and LEF (around 20% of MC) in BSC is suggested to produce NADPH for reduction of oxaloacetate (Höfer et al., 1992; Meister et al., 1996). Furthermore, some of the 3-phosphoglycerate produced by RuBisCO carboxylation are exported to the MC and converted to triosephosphate, and ATP and NADPH are consumed in this step, as observed by the high activity of phosphoglyceratekinase and NADP-triosephosphate dehydrogenase in both MC and BSC in many C_4_ plants, including NADP-ME types (Ku and Edwards, 1975; Kanai and Edwards, 1999). Taking this metabolism into account, if 50% of 3-phosphoglycerate is exported and converted to triosephosphate in MC, estimated ATP/NADPH demand is much higher in BSC (ATP/NADPH = 5) than in MC (ATP/NADPH = 1.9) in C_4_ *Flaveria* (Munekage and Taniguchi, 2016). Leakage of CO_2_ from BSC to MC also should be considered for evaluating energy costs. Assuming that 20% of the concentrated CO_2_ leaked from BSC to MC (Henderson et al., 1992), the ATP/NADPH demand would increase to 2 in MC and 8 in BSC in C_4_ *Flaveria*. Since NDH subunits are three times more abundant in BSCs than in MCs in *F. bidentis* (Nakamura et al., 2013) and ATP demand is very high in BSC, we speculate that the suppression of NDH-dependent CEF activity limits metabolism, especially in BSCs. On the other hand, suppression of NDH-dependent CEF activity did not impair ΔpH generation in MCs, considering that NPQ and ECSt were not lowered in *NdhO*-RNAi plants compared with those in WT plants (Fig. 4, C and D). Taken together, suppression of NDH-dependent CEF activity may limit Calvin-Benson cycle activity in BSCs and simultaneously affect coordinated metabolism, including the C_4_ cycle and part of the Calvin-Benson cycle (the conversion step of 3-phosphoglycerate to triosephosphate) taking place in MCs, thereby decreasing ATP consumption in chloroplasts of MCs. Thus, the high NPQ and ECSt in the MCs of *NdhO*-RNAi plants may be attributed to excess ΔpH resulting from reduced ATP consumption and residual PGR5/PGRL1-dependent CEF activity.

In conclusion, the NDH-dependent route contributes to the C_4_ photosynthesis as a major route of CEF to provide ATP in the chloroplast. The PGR5/PGRL1-dependent route also partly contributes to C_4_ photosynthesis and is vital for NPQ induction at high light in *F. bidentis*.

## Materials and Methods

### Plant materials and growth conditions

*Flaveria bidentis* plants were grown in pots filled with soil and vermiculite (3:2) in a growth chamber set at 24 °C with a 12-h light/12-h dark photoperiod (250 μmol photons m^-2^ s^-1^). The WT, vector control, *PGR5*-RNAi, and *PGRL1*-RNAi plants were grown for 8–10 weeks, whereas the *NdhO*-RNAi plants were grown for 12–16 weeks. The fourth-seventh mature leaves were used in the subsequent experiments.

### RNAi construct preparation and transformation of Flaveria bidentis

The *PGR5, PGRL1*, and *NdhO* target sequences (Supplemental Table S1) were amplified from cDNA derived from the WT *F. bidentis* and inserted into the pART7 vector in sense and antisense orientations to generate hairpin RNA sequences under the control of the cauliflower mosaic virus (CaMV) 35S promoter. These cassettes containing the CaMV 35S promoter and the *ocs* terminator were subcloned into the pART27 binary vector at the *Not*I restriction enzyme site (Supplemental Fig. S6). The resulting RNAi vectors were introduced into *F. bidentis* via *Agrobacterium tumefaciens* strain AGL1 (Chitty et al., 1994). Transformants were selected based on kanamycin resistance.

### Quantitative real-time PCR

Total RNA was extracted from leaves with the RNAs-ici!-P kit (RIZO Inc., Tsukuba, Japan). The extracted total RNA was then treated with the RNase-Free DNase Set (Qiagen, Venlo, the Netherlands) on a column, purified with the NucleoSpin RNA Clean-up XS (Macherey-Nagel GmbH & Co. KG, Düren, Germany) and then used as the template for reverse transcription with the ReverTra Ace qPCR RT kit (Toyobo, Osaka, Japan). Quantitative real-time PCR assays were completed with the SYBR Green Master Mix (Thermo Fisher Scientific, Waltham, USA) and the LightCycler 96 system (Roche, Basel, Switzerland). Details regarding the quantitative real-time PCR primers are provided in Supplemental Table S2. The target transcript levels were quantified relative to the expression level of actin7 (*ACT7*), which was used as the reference gene.

### Immunoblot analysis

Leaf samples were crushed in liquid nitrogen, and the resultant powder was suspended in an extraction buffer comprised of 50 mM Tris-HCl (pH 8.0) and 1% Protease Inhibitor Cocktail (Sigma-Aldrich), followed by centrifugation at 20,400 × *g* for 10 min at 4 °C. The supernatant was used as the soluble protein sample. The pellet was resuspended in the extraction buffer containing 2% SDS, followed by incubation at 37 °C for 20 min. After centrifugation at 15,000 rpm for 10 min at room temperature, the supernatant was used as the membrane protein sample. For total protein extraction, leaf powder was suspended in an extraction buffer containing 2% SDS and centrifuged at 15,000 rpm for 10 min at 20 °C. Protein samples were mixed with 4× Laemmli Sample Buffer (Bio-Rad) containing 10% 2-mercaptoethanol, denatured at 95 °C for 5 min, and separated by 12% or 15% sodium dodecyl sulfate-polyacrylamide gel electrophoresis. The separated proteins were transferred to polyvinylidene difluoride membranes (Immobilon-P; Merck Millipore, Burlington, MA, USA) and analyzed with the following antibodies: anti-NdhH antibodies (provided by Gilles Peltier), anti-PGR5 antibodies (Munekage et al., 2002), anti-PGRL1 antibodies (raised from recombinant proteins provided by Toru Hisabori), anti-Rieske antibodies (Sanda et al., 2011), anti-PsbO antibodies (provided by the late Akira Watanabe), anti-PsaD antibodies (purchased from Agrisera, Vännäs, Sweden), anti-PEPC antibodies (provided by Tsuyoshi Furumoto), and anti-RbcL antibodies (provided by Hiroki Ashida). Immunocomplexes were detected with the Pierce ECL Plus Western Blotting Substrate (Thermo Fisher Scientific). Chemifluorescent signals were detected with the ImageQuant LAS-4000 mini Lumino image analyzer (Fujifilm, Tokyo, Japan).

### Isolation of ruptured chloroplasts

Leaves were homogenized at high speed (5,000–10,000 rpm) using a Polytron homogenizer PT10-35GT (Kinematica AG, Switzerland) to grind MCs and BSCs in a medium comprising 330 mM sorbitol, 50 mM Tricine-KOH (pH 8.4), 5 mM MgCl_2_, 10 mM NaCl, and 2 mM ascorbate. The solution was centrifuged at 3,000 × *g* for 2 min at 4 °C. The supernatant was used to evaluate the content of RbcL and PEPC. The pellet was suspended in a medium consisting of 330 mM sorbitol, 20 mM HEPES-NaOH (pH 7.6), 5 mM MgCl_2_, and 2.5 mM EDTA and then centrifuged at 3,000 × *g* for 2 min at 4 °C. The pellet, which contained the thylakoid membrane, was suspended in a specific medium, as described by Munekage et al. (2002). The suspension was adjusted to a concentration of 20 μg chlorophyll ml^-1^ prior to measuring chlorophyll fluorescence.

### Chlorophyll fluorescence measurement

Chlorophyll fluorescence was measured with the MINI-PAM pulse-amplitude fluorometer (Walz, Effeltrich, Germany) equipped with a light-emitting diode (emission maximum at 650 nm) as the measuring light source and a halogen lamp (filtered to provide a wavelength < 710 nm) as the actinic light and a saturating pulse source. The ferredoxin-dependent reduction of PQ in ruptured chloroplasts was measured under illumination provided by a weak measuring light (0.25 μmol photons m^-2^ s^-1^) as described previously (Endo et al., 1998). NADPH (250 μM) and spinach ferredoxin (5 μM) were used as electron donors. In this assay, electrons are transferred from NADPH to ferredoxin by reverse reaction of ferredoxin-NADP^+^ oxidoreductase and further transferred to PQ by NDH-complex or PGR5/PGRL1-dependent cyclic activity (Munekage et al., 2004; DalCorso et al., 2008). Time-dependent chlorophyll fluorescence level (Ft) was normalized by setting the minimal fluorescence (Fo)before electron donor addition and the maximal fluorescence (Fm) during saturating pulse irradiation to 0 and 1, respectively. The *in vitro* assay of the linear electron transport activity was completed in the presence of an electron acceptor (25 μM methyl viologen) as described by Munekage et al. (2002). Moreover, Fo (minimal fluorescence in darkness), Fm (maximal fluorescence in darkness), Fs (minimal fluorescence under illumination provided by actinic light), and Fm′ (maximal fluorescence under illumination provided by actinic light) were measured for the subsequent calculation of Fv/Fm = (Fm - Fo)/Fm (Kitajima and Butler, 1975) and Φ_PSII_ = (Fm′ – Fs)/Fm′ (Genty et al., 1989).

### Simultaneous measurements of CO_2_ exchange and chlorophyll fluorescence

The leaf CO_2_ assimilation rate was measured with the LI-6400XT gas analyzer equipped with the 6400-40 Leaf Chamber Fluorometer (LI-COR, Lincoln, NE, USA), as described by Munekage et al. (2008). The CO_2_ gas exchange occurred at 25 °C and 50% relative humidity. 90% red LEDs peaking at 635 nm and 10% blue LEDs peaking at 470 nm were used for actinic light. Plants were not dark-adapted and were immediately transferred from the growth chamber to the gas analyzer for measurement. To measure the response to light intensity, we performed the experiments under low to high light intensity. For measurements of response to intercellular CO_2_ concentration, experiments began with an ambient CO_2_ concentration of 400 ppm followed by an increase or decrease in CO_2_ concentration. The initial slope of the intercellular CO_2_ concentration-response curve was calculated from the slope of the regression line of the three data points at ambient CO_2_ concentrations below 100 ppm. Additionally, chlorophyll fluorescence was simultaneously measured, with rETR and NPQ calculated as follows: rETR =Φ_PSII_ × PAR (photosynthetically active radiation) and NPQ = (Fm – Fm′)/Fm′ (Bilger and Björkman, 1990), where rETR does not account for changes in leaf absorbance or partitioning of light between photosystems. Dark-acclimation for 15 min was performed for measurements of dark-acclimated state but not for those of light-acclimated state.

### Spectroscopic measurement of the electrochromic shift

The ECS signal was monitored based on the changes in absorbance at 515 nm detected with the Dual-PAM 100 measuring system equipped with a P515/535 module (Walz) as described by Nishikawa et al. (2012). The ECSt was monitored according to the amplitude of the decline in the ECS signal after turning off the actinic light source (100, 330, or 1130 μmol photons m^-2^ s^-1^ for 4 min, respectively). A representative trace of the ECS signal after turning off the actinic light is presented in Supplemental Fig. S7. The ECSt level was normalized based on the changes in absorbance at 515 nm induced by a single turnover saturating flash.

### Estimation of the chlorophyll content per leaf area

Chlorophyll was extracted from a crushed leaf disc with 80% aqueous acetone. The absorbance of the chlorophyll extract in a 10-mm cuvette was measured with the NanoDrop 2000c spectrophotometer (Thermo Fisher Scientific). The chlorophyll concentration of the extracts in 80% acetone was determined, as described by Porra et al. (1989).

### Staining of leaves with toluidine blue

Leaves were fixed in a buffer comprising 2% paraformaldehyde, 50% ethanol, 1 mM CaCl_2_, and 50 mM PIPES-NaOH (pH 7.2). The sample was embedded in Paraplast X-TRA (melting point of 50–54 °C; Sigma-Aldrich, USA), after which 6-μm sections were prepared with a microtome. The sections were stained with 0.02% toluidine blue for 3 min.

### Accession numbers

The *PGR5A, PGR5B*, and *PGR5C* cDNA sequences were submitted to the DDBJ (LC493040, LC493041, and LC493042, respectively).

## Acknowledgments

This work was supported by Next Generation World-Leading Research (Grant No. GS019) and JSPS KAKENHI (Grant No. 16H06557). We thank Kaoru Morikawa, Risa Kishizaki and Kikuko Sumiya for technical assistance.

## Author contributions

Y. N. M. designed the study; T. O., K. K., N. N. and Y. Y. T. performed the experiments; T. S. helped with ECS measurement; A. Y. participated in helpful discussions; T. O. and Y. N. M. wrote the manuscript.

## Funding information

This work was supported by Next Generation World-Leading Research (Grant No. GS019) and JSPS KAKENHI (Grant No. 16H06557).

## Supplemental Data

**Supplemental Figure S1.** Immunoblot images used to quantify the relative content of membrane proteins. Quantitative values shown in Figure 1C were calculated from the chemiluminescence signal intensities of these samples and the samples shown in Figure 1B. Lanes were loaded with 20 μg protein to detect PGR5 and 10 μg to detect PGRL1, NdhH, Rieske, PsaD, and PsbO.

**Supplemental Figure S2.** Evaluation of the relative content of MC- and BSC-specific proteins in thylakoid extracts used to measure CEF activity. The amounts of PEPC localized to MCs and RbcL localized to BSCs, in soluble proteins of thylakoid extracts from wild-type and each transgenic line were compared to those in soluble proteins of wild-type leaves. Lanes were loaded with 5 μg protein for immune detection of PEPC and RbcL, and for Coomassie blue staining.

**Supplemental Figure S3.** Growth of the wild-type (WT), vector control (VC), *PGR5*-RNAi #3, *PGRL1*- RNAi #4, and *NdhO*-RNAi #1, #7, and #18 plants under 1,000 μmol photons m^-2^ s^-1^ illumination for 35 days. (A) Observable phenotypes of plants. The bar indicates 5 cm. (B) Leaf area per plant. Vertical bars indicate the SD (n = 5). Asterisks indicate significant differences (Student *t*-test, *P* < 0.05) between WT and *PGR5*-RNAi, *PGRL1*-RNAi, or *NdhO*-RNAi lines.

**Supplemental Figure S4.** Relationship between the net CO_2_ assimilation rate and rETR measured at 2,000 μmol photons m^-2^ s^-1^ (see also Fig. 4, A and C). Black closed circle, WT; black open circle, vector control; light blue square, *PGR5*-RNAi #3; blue diamond, *PGRL1*-RNAi #4; red triangle, *NdhO*-RNAi #1; pink inverted triangle, *NdhO*-RNAi #18. Vertical and parallel bars indicate the SD (n = 3–6).

**Supplemental Figure S5.** The amounts of PEPC and RbcL in wild-type, vector control, *PGR5*-RNAi #3, *PGRL1*-RNAi #4, and *NdhO*-RNAi #1, #7, and #18 plants. (A) Immunoblot analysis of PEPC and RbcL and Coomassie blue staining of loaded proteins performed on samples extracted from three different plants. Lanes were loaded with 0.2 μg chlorophyll for immune detection of PEPC and RbcL, or 0.5 μg chlorophyll for Coomassie blue staining. (B) Relative content of PEPC or RbcL quantified by chemiluminescence signal intensities of immunoblot analysis. The signal intensity of the WT plants was set as 1. Vertical bars indicate the SD (n=3).

**Supplemental Figure S6.** RNA interference constructs in the pART27 binary vector targeting *FbPGR5* (A), *FbPGRL1* (B), and *FbNdhO* (C), as well as the vector control (D). CaMV 35S, cauliflower mosaic virus 35S promoter; PPDK first intron, first intron of the pyruvate phosphate dikinase gene; ocsT, octopine synthase terminator; nosP, nopaline synthase (nos) promoter; npt II, neomycin phosphotransferase gene; nosT, nos terminator; RB, the right border of T-DNA; LB, left border of T-DNA.

**Supplemental Figure S7.** Representative trace of the ECS signal to determine the ECSt parameter. The ECSt level was determined as the amplitude of the decline in the ECS signal after turning off the actinic light illuminated for 5 min, and was normalized based on the changes in absorbance at 515 nm induced by a single turnover saturating flash.

**Supplemental Table S1.** RNA interference target sequences

**Supplemental Table S2.** Sequences of primers used for quantitative real-time PCR

